# Homeostatic depression shows heightened sensitivity to synaptic calcium

**DOI:** 10.1101/2020.10.12.336883

**Authors:** Catherine J. Yeates, C. Andrew Frank

**Affiliations:** Department of Anatomy and Cell Biology, University of Iowa Carver College of Medicine, Iowa City, IA, USA; Interdisciplinary Graduate Program in Neuroscience, University of Iowa, Iowa City, IA, USA; Iowa Neuroscience Institute, University of Iowa Carver College of Medicine, Iowa City, IA, USA; Department of Biology, University of Dayton, Dayton, OH, USA

**Keywords:** synapse, homeostasis, depression, *Drosophila melanogaster*, NMJ, neurotransmission, plasticity

## Abstract

Synapses and circuits rely on homeostatic forms of regulation in order to transmit meaningful information. The *Drosophila melanogaster* neuromuscular junction (NMJ) is a well-studied synapse that shows robust homeostatic control of function. Most prior studies of homeostatic plasticity at the NMJ have centered on presynaptic homeostatic potentiation (PHP). PHP happens when postsynaptic muscle neurotransmitter receptors are impaired, triggering retrograde signaling that causes an increase in presynaptic neurotransmitter release. As a result, normal levels of evoked excitation are maintained. The counterpart to PHP at the NMJ is presynaptic homeostatic depression (PHD). Overexpression of the Drosophila vesicular glutamate transporter (VGlut) causes an increase in the amplitude of spontaneous events. PHD happens when the synapse responds to the challenge by decreasing quantal content during evoked neurotransmission – again, resulting in normal levels of postsynaptic excitation.

We hypothesized that there may exist a class of molecules that affects both PHP and PHD. Impairment of any such molecule could hurt a synapse’s ability to respond to any significant homeostatic challenge. We conducted an electrophysiology-based screen for blocks of PHD. We did not observe a block of PHD in the genetic conditions screened, but we did find loss-of-function conditions that led to a substantial deficit in evoked amplitude when combined with VGlut overexpression. The conditions causing this phenotype included a double heterozygous loss-of-function condition for genes encoding the inositol trisphosphate receptor (IP_3_R – *itpr*) and ryanodine receptor (*RyR*). IP_3_Rs and RyRs gate calcium release from intracellular stores. Pharmacological agents targeting IP_3_R and RyR recapitulated the genetic losses of these factors, as did lowering calcium levels from other sources. Our data are consistent with the idea that the homeostatic signaling process underlying PHD is especially sensitive to levels of calcium at the presynapse.

## 1 Introduction

Animal nervous systems use forms of homeostatic synaptic plasticity to maintain stable function. Over the last 20-25 years, studies from diverse systems have revealed a wealth of information about how forms of homeostatic synaptic plasticity are implemented (Davis, 2013; Davis and Müller, 2015; Delvendahl and Müller, 2019; Marder and Goaillard, 2006; Pozo and Goda, 2010; Turrigiano, 2008). In particular, the *Drosophila melanogaster* neuromuscular junction (NMJ) has uncovered many facets of homeostatic implementation on a molecular level (Frank, 2014a; Frank et al., 2020). Much of the NMJ homeostasis work in both Drosophila and vertebrates has focused on a form of homeostatic plasticity termed presynaptic homeostatic potentiation (PHP). With PHP, manipulations that impair postsynaptic muscle receptor function trigger an increase in presynaptic vesicle release (Cull-Candy et al., 1980; Davis et al., 1998; Frank et al., 2006; Petersen et al., 1997; Wang et al., 2016).

Homeostatic plasticity at the NMJ is a bi-directional process. First, PHP is reversible – when manipulations that impair muscle receptor function are removed, the presynaptic potentiation ceases (Wang et al., 2016; Yeates et al., 2017). Second, the Drosophila NMJ can depress quantal content in a homeostatic manner functionally opposite to PHP: Presynaptic homeostatic depression (PHD). Experimentally, one way to trigger PHD is to overexpress the *Drosophila* vesicular glutamate transporter gene, *VGlut*, in motor neurons. Overexpression of the glutamate transporter leads to an increase in the diameter of glutamatergic vesicles, an increase in quantal size across the entire distribution of spontaneous miniature events, and very large spontaneous quantal events (Daniels, 2004). To compensate for this, quantal content at the NMJ is lowered, resulting in normal evoked postsynaptic excitation (Daniels et al., 2004).

Many genes have been shown to be necessary for PHP at the NMJ. But much less is known about PHD. PHP and PHD result in opposite changes in quantal content, and studies suggest divergent and separable mechanisms governing these forms of homeostatic plasticity. Some genes required for homeostatic potentiation are dispensable for homeostatic depression (Gaviño et al., 2015; Li et al., 2018; Marie et al., 2010). Moreover, unlike homeostatic potentiation, homeostatic depression does not appear to involve a change in the size of the readily releasable pool of synaptic vesicles (Li et al., 2018). Rather, homeostatic depression appears to involve a decrease in release probability (Gaviño et al., 2015). Finally, PHP at the NMJ appears to be a process that is dependent on the input (i.e. the type of synapse formed at the NMJ) (Newman et al., 2017) while PHD does not appear to be input specific (Li et al., 2018).

The degree of overlap between homeostatic depression and homeostatic potentiation is unknown. We designed a small-scale, directed screen to test for links between these two forms of homeostatic plasticity. For the screen, we targeted genes based on prior evidence that their impairment in the neuron caused a failure of the long-term maintenance of PHP. We examined loss-of-function conditions for these genes in a VGlut overexpression background for PHD. We did not find any cases of failed homeostatic depression – the conditions we examined showed decreases in quantal content in response to increased quantal size. However, we did find an interesting and unexpected evoked neurotransmission phenotype: a robust decrease in excitatory postsynaptic potential (EPSP) amplitude in a VGlut overexpression background. We observed this phenotype for a double heterozygous loss-of-function condition for the Ryanodine and IP_3_ receptors. In our follow-up work, pharmacology phenocopied this genetic result, and our overall findings are consistent with the idea that the PHD system may show a heightened sensitivity to low calcium.

Our findings highlight a novel synaptic transmission phenotype. Prior characterizations of homeostatic depression do not report decreases in EPSP amplitude in VGlut overexpression relative to controls (Daniels et al., 2004; Gaviño et al., 2015; Li et al., 2018; Marie et al., 2010). Studies at the NMJ generally suggest models in which homeostatic compensation maintains evoked neurotransmission at the synapse approximately at control levels (Davis, 2013). Our results suggest that impairing store calcium channels may result in a cumulative defect in neurotransmission when there is a concurrent PHD challenge. We find this interesting, especially in light of the fact that these same store channels are required for the maintenance of PHP (James et al., 2019) and because other recent studies in other systems have implicated store calcium in presynaptic release mechanisms (e.g., (de Juan-Sanz et al., 2017)).

## 2 Materials and Methods

### Drosophila stocks and husbandry

Fruit fly stocks were obtained from the Bloomington Drosophila Stock Center (BDSC, Bloomington, Indiana), Kyoto Stock Center (DGRC, Kyoto, Japan), Japan National Institute of Genetics (Mishima, Shizuoka, Japan), Vienna Drosophila Research Center (VDRC, Vienna, Austria), or from the labs that generated them. *w*^*1118*^ was used as a wild-type (WT) control (Hazelrigg et al., 1984). RNAi lines and mutants used in the screen are reported in Supplemental Table S1.

Fruit flies were raised on cornmeal, molasses, and yeast medium (see BDSC website for standard recipe) in temperature-controlled conditions. Animals were reared at 25°C until they reached the wandering third instar larval stage, at which point they were selected for electrophysiological recording. *UAS-VGlut* (Daniels et al., 2004) was recombined with *OK371-GAL4* (Mahr and Aberle, 2006; Meyer and Aberle, 2006) to drive constitutive overexpression of VGlut. The full genotype of these animals is: *w; VGlut, OK371-Gal4/CyO-GFP*. Virgins of these flies were crossed to RNAi lines or mutants to test for changes to homeostatic depression. *w; OK371-Gal4/+* was used as a genetic control for baseline electrophysiology.

### Electrophysiology and analysis

Larvae were dissected in a modified HL3 saline comprised of: NaCl (70 mM), KCl (5 mM), MgCl_2_ (10 mM), NaHCO_3_ (10 mM), sucrose (115 mM = 3.9%), trehalose (4.2 mM = 0.16%), HEPES (5.0 mM = 0.12%), and CaCl_2_ (0.5 mM, except as noted).

For pharmacology, Dantrolene (R&D Systems) and Xestospongin C (Abcam) were used. Dantrolene was mixed into saline to a final concentration of 25 µM. Larvae were cut open on the dorsal side and allowed to incubate in the Dantrolene saline for 5 minutes. The rest of the dissection and recording was completed in Dantrolene saline. Xestospongin C was applied in a similar manner, with the animals allowed to incubate in 20 µM Xestospongin C saline for 5 minutes before they were recorded, also in saline containing Xestospongin C.

Electrophysiological data were collected using an Axopatch 200B amplifier (Molecular Devices, Sunnyvale, CA) in bridge mode, digitized using a Digidata 1440A data acquisition system (Molecular Devices), and recorded with pCLAMP 10 acquisition software (Molecular Devices). A Master-8 pulse stimulator (A.M.P. Instruments, Jerusalem, Israel) and an ISO-Flex isolation unit (A.M.P. Instruments) were utilized to deliver 1 ms suprathreshold stimuli to the appropriate segmental nerve. The average spontaneous miniature excitatory postsynaptic potential (mEPSP) amplitude per NMJ was quantified by hand, approximately 100 individual spontaneous release events per NMJ (MiniAnalysis, Synaptosoft, Fort Lee, NJ). Measurements from all NMJs of a given condition were then averaged. For evoked neurotransmission, 30 excitatory postsynaptic potentials (EPSPs) were averaged to find a value for each NMJ. These were then averaged to calculate a value for each condition. Quantal content (QC) was calculated by the ratio of average EPSP and average mEPSP amplitudes for each individual NMJ. An average quantal content was then calculated for each condition. EPSP variability was assessed by measuring each of the 30 traces individually and calculating a standard deviation and range for that NMJ. Range was defined as the maximum EPSP value minus the minimum EPSP value.

### Immunostaining

An immunostaining experiment is detailed for Figure 4. Procedures match those previously published (Brusich et al., 2015; Brusich et al., 2018; James et al., 2019; Spring et al., 2016; Yeates et al., 2017). Briefly, third instar larvae were filleted and fixed for 5 minutes with Bouin’s fixative (Ricca Chemical, Arlington, TX). After washes, fixed fillets were incubated in primary antibodies overnight at 4ºC, mouse anti-Brp (nc82, 1:250, University of Iowa Developmental Studies Hybridoma Bank) (Wagh et al., 2006) and rabbit anti-Dlg (1:5,000) (Budnik et al., 1996). After washes, fillets were incubated in fluorophore-conjugated secondary antibodies overnight at 4ºC (Jackson ImmunoResearch Labs, West Grove, PA), goat anti-mouse-488 (DyLight, 1:1000) and goat anti-rabbit-549 (DyLight, 1:2000). After washes, fillets were mounted and Dlg boutons were counted blinded by hand and double checked for Brp signal in apposition.

### Statistical Analyses

Statistical analyses were conducted using GraphPad Prism Software. Statistical significance was assessed either by Student’s T-Test when one experimental data set was being directly compared to a control data set, or one-way ANOVA with Tukey’s post-hoc test when multiple data sets were being compared. Specific *p* value ranges are noted in the Figure legends and shown in graphs as follows: * *p* < 0.05, ** *p* < 0.01, and *** *p* < 0.001 (* and # are used in Figures if there are additional comparisons highlighted). For some comparisons that are close to *p* < 0.05 statistical significance but do not achieve it (0.05 < *p* < 0.1), specific values are reported on the graph itself. Calcium cooperativity data were analyzed using a non-linear fit regression analysis on GraphPad Prism.

## 3 Results

### A recombinant line to analyze presynaptic homeostatic depression (PHD)

Using previously published reagents, we generated a fly stock with constitutive *VGlut* transgene overexpression. Such a stock could be used as a tool for a single-cross genetic screen. To generate the stock, we recombined the *OK371-Gal4* motor neuron driver (Mahr and Aberle, 2006; Meyer and Aberle, 2006) with a *UAS-VGlut* transgene (Daniels et al., 2004). We placed these two genetic elements in *cis* on *Drosophila melanogaster* Chromosome II. *OK371-Gal4* is an enhancer trap line for the *VGlut* promoter itself. This ensured that GAL4-driven *UAS-VGlut* overexpression would happen in desired tissues, *Drosophila* motor neurons.

We tested if the recombinant line constitutively overexpressing *UAS-VGlut* could express PHD at the NMJ. We crossed the recombinant stock to our wild-type stock (*w*^*1118*^, herein: WT) (Cross result, herein: “*VGlut, OK371/+*”). By NMJ electrophysiology, we recorded from WT control, *OK371/+* control, and *w; VGlut, OK371/+*. As expected, *VGlut, OK371/+* NMJs showed an increase in spontaneous miniature excitatory postsynaptic potential (mEPSP) amplitude compared to controls (Fig. 1A-C; data also in Supplementary Table S1). Compared to WT control NMJs, there was no significant difference in evoked postsynaptic amplitudes for *VGlut, OK371/+* NMJs (Fig. 1D; *p = 0*.*82*, one-way ANOVA). This was because of an accurate homeostatic decrease in quantal content (QC) (Fig 1E) – hence, successful PHD. This result matched prior studies that had used WT as a control and a *trans OK371/UAS-VGlut* combination to induce PHD (Daniels et al., 2004; Gaviño et al., 2015; Li et al., 2018).

**Figure 1.**
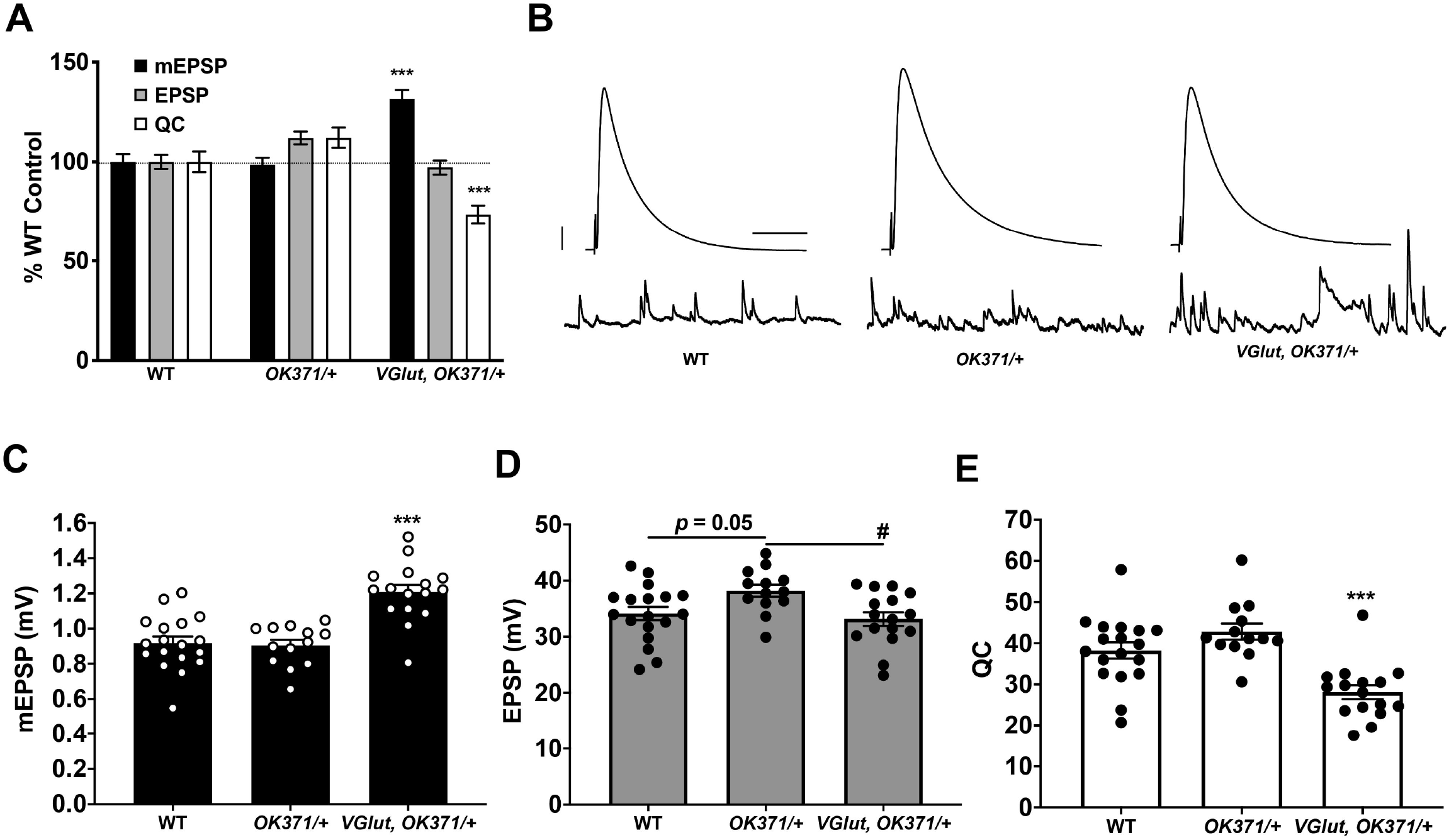
Presynaptic homeostatic depression (PHD) works successfully with a recombinant line of *OK371-Gal4* and *UAS-VGlut*. **(A)** NMJ electrophysiological data for miniature excitatory postsynaptic potentials (mEPSP), excitatory postsynaptic potentials (EPSP), and quantal content (QC). Data are normalized to WT (*w*^*1118*^) values. *VGlut, OK371/+* NMJs have increased mEPSP but normal EPSP because of decreased QC, indicative of successful PHD. (*** *p < 0*.*001* vs. WT by one-way ANOVA with Tukey’s post-hoc). **(B)** Representative electrophysiological traces. Large traces are EPSPs; small traces are mEPSPs. Scale bars for EPSPs (mEPSPs) are 5 mV (1 mV) and 50 ms (1000 ms). **(C)** Raw data for mEPSPs. **(D)** Raw data for EPSPs. **(E)** Raw data for QC. For (C)-(E), bars are averages and error bars are ± SEM. *** *p < 0*.*001* vs. WT or vs. *OK371/+*; # *p < 0*.*05* vs. *OK371/+*; analyses by one-way ANOVA with Tukey’s post-hoc.

Even though PHD was successful relative to WT for our test cross, we noted a small, but statistically significant, baseline increase in the EPSP amplitude of *OK371/+* NMJs. This increase in *OK371/+* EPSP level was present compared either to WT control or to *VGlut, OK371/+* (Fig. 1D). One possibility is that the *OK371/+* genetic background has slightly elevated release, and the combined addition of *UAS-VGlut* reveals a slight depression in evoked amplitude. Noting this potentially important difference in our driver control, we continued using the *OK371/+* heterozygous condition as a genetic background control. *OK371/+* is a closer genetic control for PHD analysis than WT.

### A genetic screen identifies an interaction between calcium stores and a PHD-inducing challenge

We used our recombinant line to conduct a genetic screen for conditions that affect presynaptic homeostatic depression (PHD). We crossed this stock to screen stocks: 1) either to drive *UAS-RNAi* transgenes to knock down genes; 2) to drive other chosen *UAS* transgenes; or 3) to combine with \ heterozygous loss-of-function mutant lines (Materials and Methods, Fig. 2A). For the screen, we targeted a subset of genes previously identified as in the neuron for homeostatic potentiation, or closely related genes. We tested 43 genotypes (sometimes multiple conditions for a single gene), including our homeostatic depression condition, *VGlut, OK371/+* (Fig. 2B, C).

**Figure 2.**
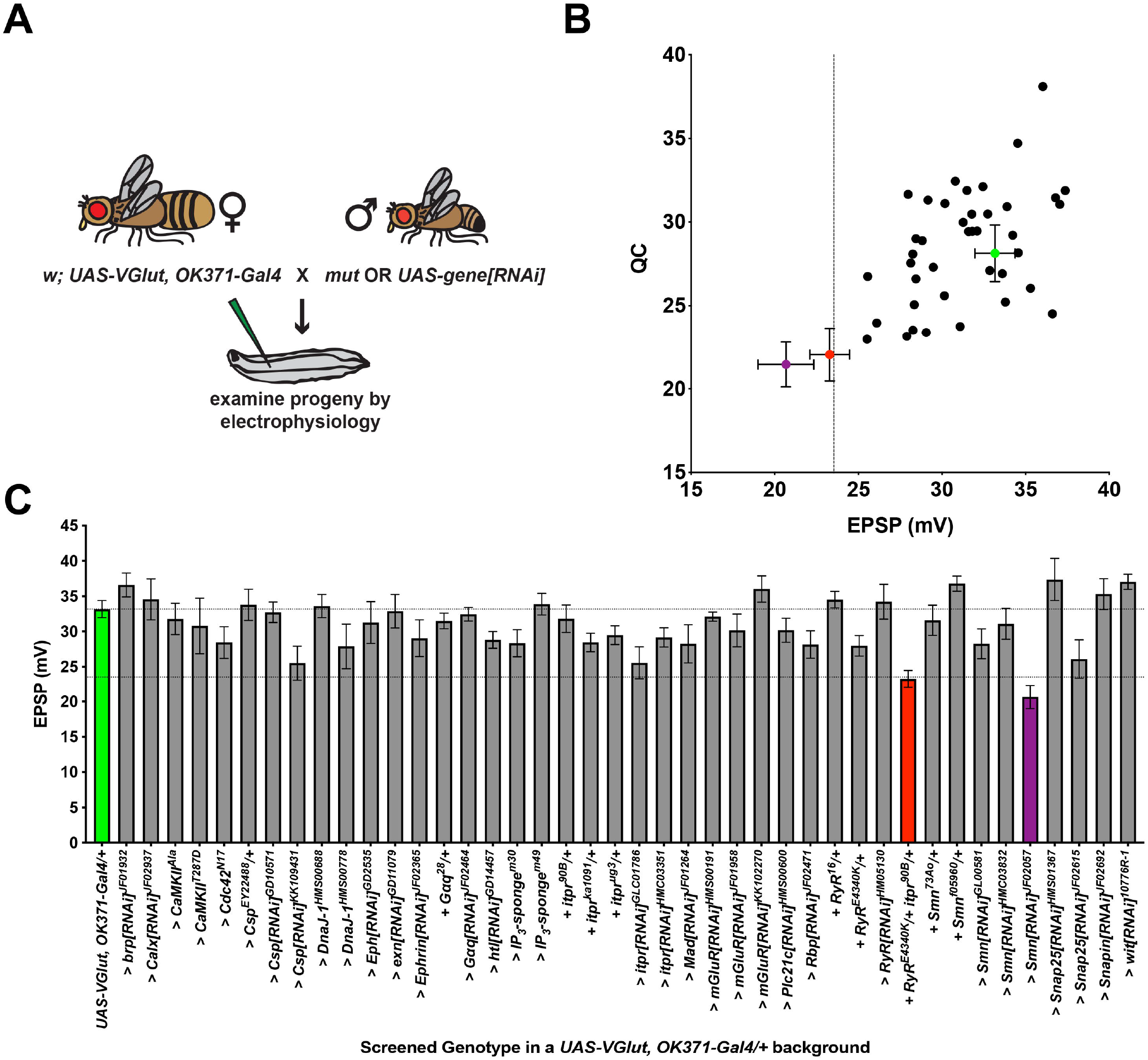
An electrophysiology screen in a PHD-challenged genetic background. **(A)** Crossing scheme for generating larvae for electrophysiological recording. Each animal recorded had a homeostatic challenge provided by VGlut overexpression and a concurrent heterozygous or RNAi condition. **(B)** Data distribution for screened conditions (x-axis = average EPSP for condition; y-axis = average QC for condition). Green = *UAS-VGlut, OK371-Gal4/+*. Red = *UAS-VGlut, OK371-Gal4/RyRE4340K; itpr90B/+*. Purple = *UAS-VGlut, OK371-Gal4/+; UAS-Smn[RNAi]*^*JF02057*^*/+*. Dotted line: EPSP value two standard deviations below *UAS-VGlut, OK371-Gal4/+* chosen as a cut off for potential follow-up hits. **(C)** Average EPSPs for screened conditions. All conditions have a *UAS-VGlut, OK371-Gal4/+* genetic background. “>“ denotes as *UAS* construct or RNAi line being driven in motor neurons by *OK371-Gal4*. “+” denotes additional mutations present as heterozygotes. Top dotted line denotes *UAS-VGlut, OK371-Gal4/+* average. Bottom dotted line denotes two standard deviations below *UAS-VGlut, OK371-Gal4/+* average.

The aggregate results of the screen are reported here (Fig. 2B, C; raw data in Supplementary Table S1). We recorded from 42 experimental heterozygous *mutant/+* or > *UAS-RNAi or -transgene/+* conditions, in the *VGlut, OK371/+* genetic background. Of those 42, 12 achieved EPSPs that were numerically larger than *VGlut, OK371/+*, and 22 achieved QCs that were numerically larger than *VGlut, OK371/+* (Fig. 2B, C). Increased evoked potentials could signify failed PHP – however, none of these cases represented statistically significant increases compared to *VGlut, OK371/+*. None were so much bigger that they were good candidates for “failed PHD.” Indeed, all of the candidates had average EPSP and QC levels below *OK371/+* NMJ baseline recordings (Compare Figs 1D, E and Fig. 2B, C).

We noted a phenotype distinct from what we were initially seeking: two crosses yielded larvae with striking decreases in NMJ EPSP amplitudes, more than two standard deviations below the average EPSPs from the baseline *VGlut, OK371/+* data set (Fig. 2B). One case was knockdown of the *Survival motor neuron* (*Smn*) gene with the *UAS-Smn[RNAi]*^*JF02057*^ line in the *VGlut, OK371/+* background. This was intriguing because Drosophila *Smn* is homologous to human *SMN*. Defects in *SMN* cause Spinal Muscular Atrophy (Lefebvre et al., 1995). Drosophila Smn has been characterized as a potential model for Spinal Muscular Atrophy (Raimer et al., 2020; Sen et al., 2011; Spring et al., 2019). Smn has also previously been implicated in PHP (Sen et al., 2011). However, the result for *UAS-Smn[RNAi]*^*JF02057*^ was not replicated by other *Smn* knockdown or loss-of-function mutant test crosses (Fig. 2B, C). We did not follow up on *Smn* for this study.

A second case with a striking decrease in EPSP amplitude in the screen was a double heterozygous genetic condition in genes encoding the Drosophila Ryanodine receptor (*RyR*) and inositol 1,4,5-trisphosphate (IP_3)_ receptor (*itpr*): *VGlut, OK371/RyR*^*E4340K*^; *itpr*^*90B*^*/+* (Fig. 2B, C). Ryanodine receptors (RyRs) and IP_3_ receptors (IP_3_Rs) are localized to the endoplasmic reticulum. They mediate release of calcium from intracellular stores (Berridge, 1984, 1987, 1998; Simkus and Stricker, 2002). The *RyR*^*E4340K*^ mutation is a single amino acid substitution (glutamic acid to lysine) (Dockendorff et al., 2000), and the *itpr*^*90B*^ mutation is a null mutant generated by imprecise excision of a transposon (Venkatesh and Hasan, 1997). We previously defined roles for RyR, IP_3_R, IP_3_ signaling and upstream components in maintaining presynaptic homeostatic potentiation (PHP) (Brusich et al., 2015; James et al., 2019).

In parallel, we screened single mutant manipulations for both genes. Neither the *RyR*^*E4340K*^*/+* heterozygous condition, nor the *itpr*^*90B*^*/+* heterozygous condition – nor any single heterozygous or RNAi knockdown condition for either gene – yielded as significantly depressed EPSPs in response to PHD challenge (Fig. 2B, C). Therefore, the screen result with the double heterozygote could be due to a genetic interaction, or it could be due to other factors in the genetic background. This preliminary finding required further characterization.

We tested if the electrophysiological phenotype could be due to a baseline neurotransmission defect when both genes are heterozygous. By electrophysiology, we compared NMJs from *OK371/RyR*^*E4340K*^; *itpr*^*90B*^*/+* larvae as a baseline double heterozygous condition vs. NMJs from *VGlut, OK371/RyR*^*E4340K*^; *itpr*^*90B*^*/+* larvae (Fig. 3A-D). Just like WT, the baseline double heterozygous condition did have a slight decrease in EPSP amplitude compared to *OK371/+* driver control (Fig. 3A). This indicated a small, but discernible defect in neurotransmission in animals where the IP_3_Rs and RyRs are both impaired. The double heterozygous condition with concurrent *VGlut* gene overexpression showed a further decrease in transmission – compared to its own genetic control, it had increased quantal size (Fig. 3B), but significantly decreased evoked amplitude (Fig. 3C) because of a large decrease in quantal content (Fig. 3D). Finally, the quantal content for *VGlut, OK371/RyR*^*E4340K*^; *itpr*^*90B*^*/+* NMJs was numerically smaller than for *VGlut, OK371/+* NMJs (Fig. 3D), but this latter numerical difference was not statistically significant (*p = 0*.*07*, one-way ANOVA).

**Figure 3.**
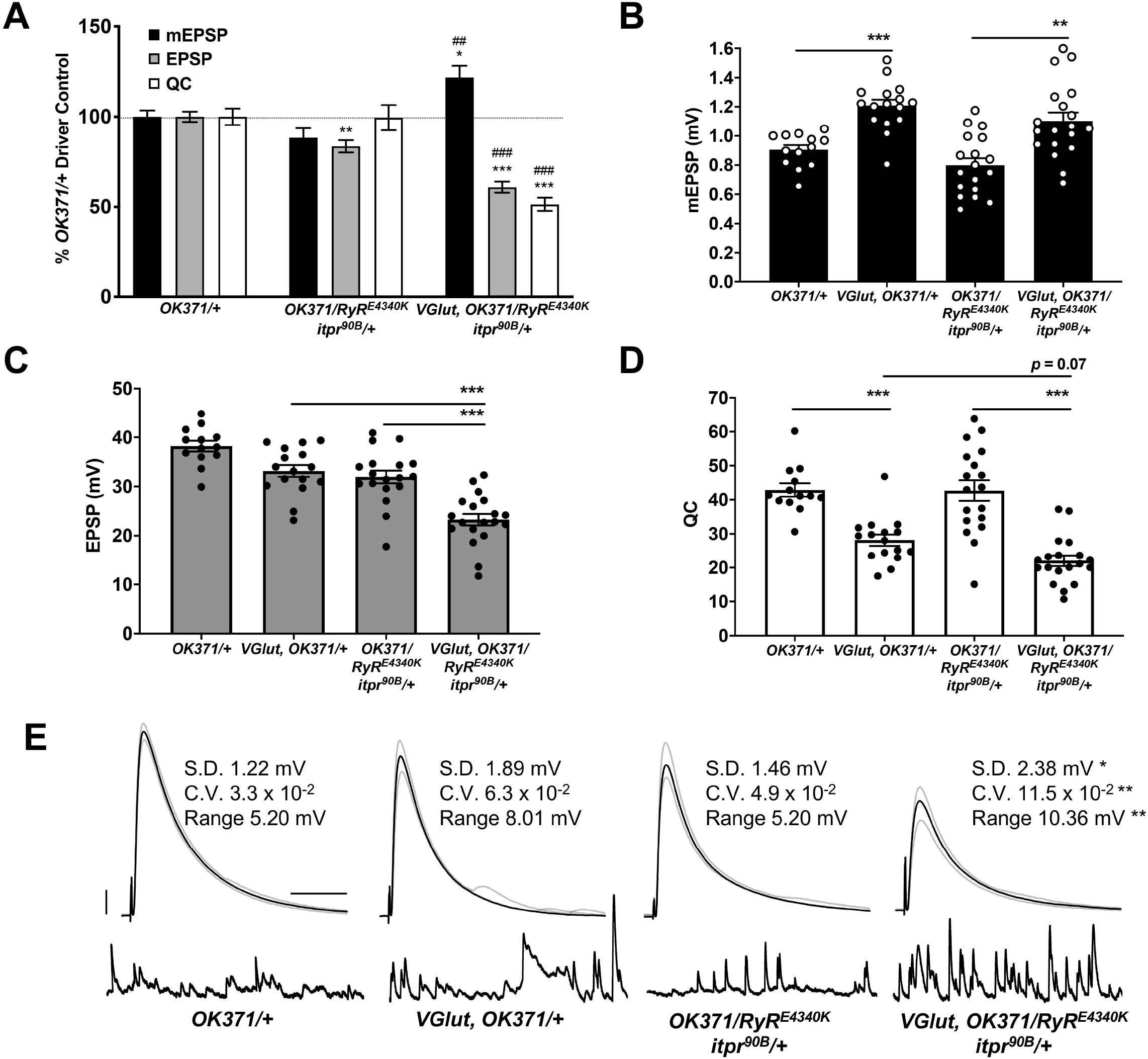
Double heterozygous loss of the *itpr* and *RyR* genes interacts with the PHD challenge to diminish neurotransmission. Note: Traces and data for *OK371/+* and *VGlut, OK371/+* are repeated from Figure 1, for genetic background comparison. Abbreviations are as in Figure 1. **(A)** NMJ electrophysiological data for mEPSP, EPSP, and QC. Data are normalized to *OK371/+* values. * *p < 0*.*05, ** p < 0*.*01*, and *\*\*\* p < 0*.*001* vs. *OK371/+*; ## *p < 0*.*01* and *### p < 0*.*001* vs. *OK371/RyRE4340K; itpr90B/+*; analyses by one-way ANOVA with Tukey’s post-hoc. **(B)** Raw data for mEPSPs. **(C)** Raw data for EPSPs. **(D)** Raw data for QC. For (B)-(D), bars are averages and error bars are ± SEM. * *p < 0*.*05, ** p < 0*.*01*, and *\*\*\* p < 0*.*001* by one-way ANOVA across genotypes, with Tukey’s post-hoc. **(E)** Representative electrophysiological traces with standard deviation (S.D.), coefficient of variation (C.V.) and range values for EPSPs. The S.D., C.V., and range were significantly higher for *VGlut, OK371/RyR*^*E4340K*^; *itpr*^*90B*^*/+* vs. its genetic control, *OK371/RyR*^*E4340K*^; *itpr*^*90B*^*/+*. * *p < 0*.*05, ** p < 0*.*01* by one-way ANOVA across genotypes, with Tukey’s post-hoc. Scale bars as in Figure 1.

We noted that the EPSP amplitude in individual *VGlut, OK371/RyR*^*E4340K*^; *itpr*^*90B*^*/+* NMJ recordings varied markedly from stimulus to stimulus. High variability could indicate unstable neuronal excitability or release. To check if evoked release events were indeed more variable, we completed additional analyses. First, we extracted the amplitude of each individual EPSP event at every NMJ recorded. From these data, we calculated the EPSP standard deviation (S.D.) and coefficient of variation (C.V.) per individual NMJ. We also calculated a range for each NMJ by subtracting the maximum EPSP of the thirty from the minimum. We averaged these S.D., C.V., and range measures for each genotype, considering all of the individual EPSP recordings. For all of these EPSP parameters, *w; VGlut, OK371/RyR*^*E4340*^; *itpr*^*90B*^*/+* animals showed statistically significant higher variability compared to controls (Fig. 3E). By contrast, double heterozygous baseline *OK371/RyR*^*E4340K*^; *itpr*^*90B*^*/+* NMJs did not differ significantly from *w; OK371/+* driver control NMJs (*p > 0*.*85* for each measure, Kruskal-Wallis ANOVA), suggesting that the variability stems from *VGlut* overexpression in the mutant background (Fig. 3E). *w; VGlut, OK371/+* NMJs showed numerically higher variability than *w; OK371/+*, but this was not statistically significant (Fig. 3E, *p > 0*.*25* for each measure, Kruskal-Wallis ANOVA).

Finally, we conducted immunostaining to check if any of these electrophysiological defects might correspond with defects in synaptic growth. We assessed growth by co-staining with antibodies against the postsynaptic PSD-95 homolog, DLG (Budnik et al., 1996) and the presynaptic active zone protein, BRP (Wagh et al., 2006). We counted boutons encased by anti-DLG signal and checked that these boutons were apposed by anti-BRP signal. By this analysis, we saw no significant changes in NMJ growth: neither the PHD challenge; nor the double heterozygous loss of the *RyR/+* and *itpr/+* genes; nor combining those manipulations together yielded significant numerical differences in bouton count (*p > 0*.*90* for all comparisons, one-way ANOVA; Fig 4A, B). One caveat to these results is that we only examined these NMJs at the level of bouton count, not at the level of the abundance of specific active zone markers (as in (Böhme et al., 2019; Goel et al., 2019; Gratz et al., 2019)).

**Figure 4.**
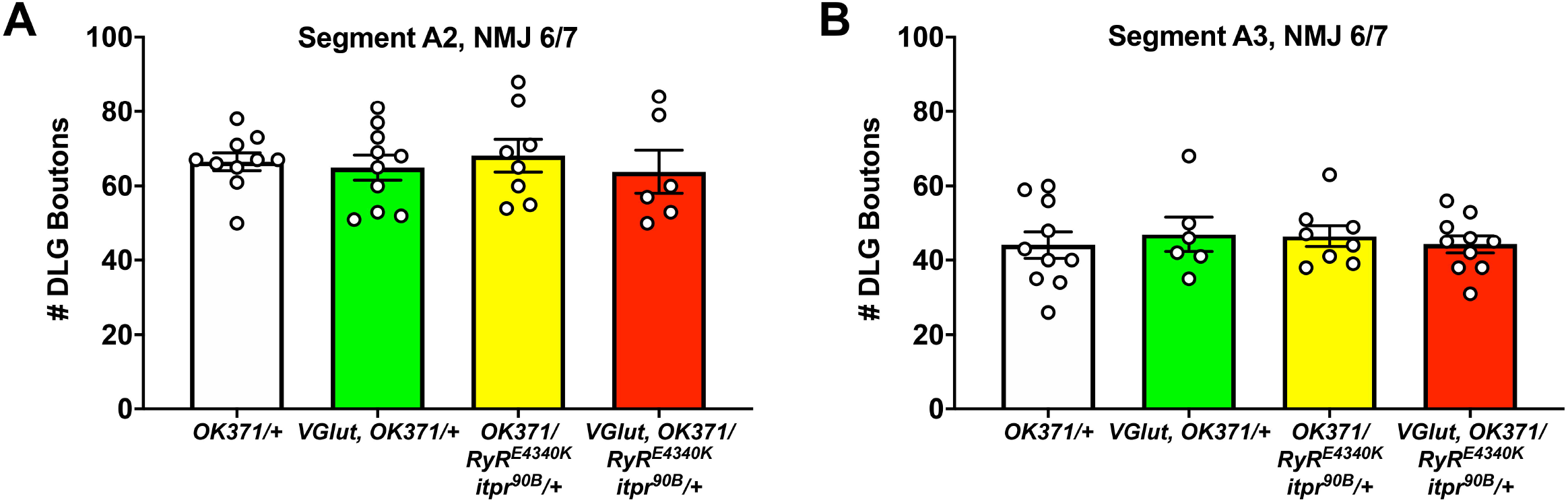
No discernible NMJ growth defects. NMJs of third instar larvae (same genotypes as Figure 3) were analysed by immunostaining, co-staining with anti-DLG for the postsynaptic density and anti-Brp to check for apposed presynaptic active zones. **(A)** DLG boutons counted for Segment A2, NMJ 6/7. **(B)** DLG boutons counted for Segment A3, NMJ 6/7. No significant differences were found across genotypes (*p > 0*.*9* for every possible head-to-head comparison, one-way ANOVA).

### Pharmacology targeting Ryanodine and IP_3_ receptors recapitulates loss-of-function genetics

We tested if the electrophysiological phenotypes we observed could be recapitulated by combining genetics and pharmacology. We started with the drug Dantrolene. Dantrolene is a RyR antagonist (Vazquez-Martinez et al., 2003; Zhao et al., 2001). In prior work at the *Drosophila* NMJ, we found that application of Dantrolene can abrogate the long-term maintenance of PHP (James et al., 2019).

We used a sensitized *OK371/+; itpr*^*90B*^*/+* genetic background. With this background, we could pharmacologically impair RyRs while also genetically impairing IP_3_Rs. We applied 25μM of Dantrolene to: 1) *OK371/+* NMJs; 2) *VGlut, OK371/+* NMJs; 3) *OK371/+; itpr*^*90B*^*/+* NMJs; and 4) *VGlut, OK371/+; itpr*^*90B*^*/+* NMJs. We also collected a set of data for genetically identical conditions without drug treatment (Fig. 5A).

**Figure 5.**
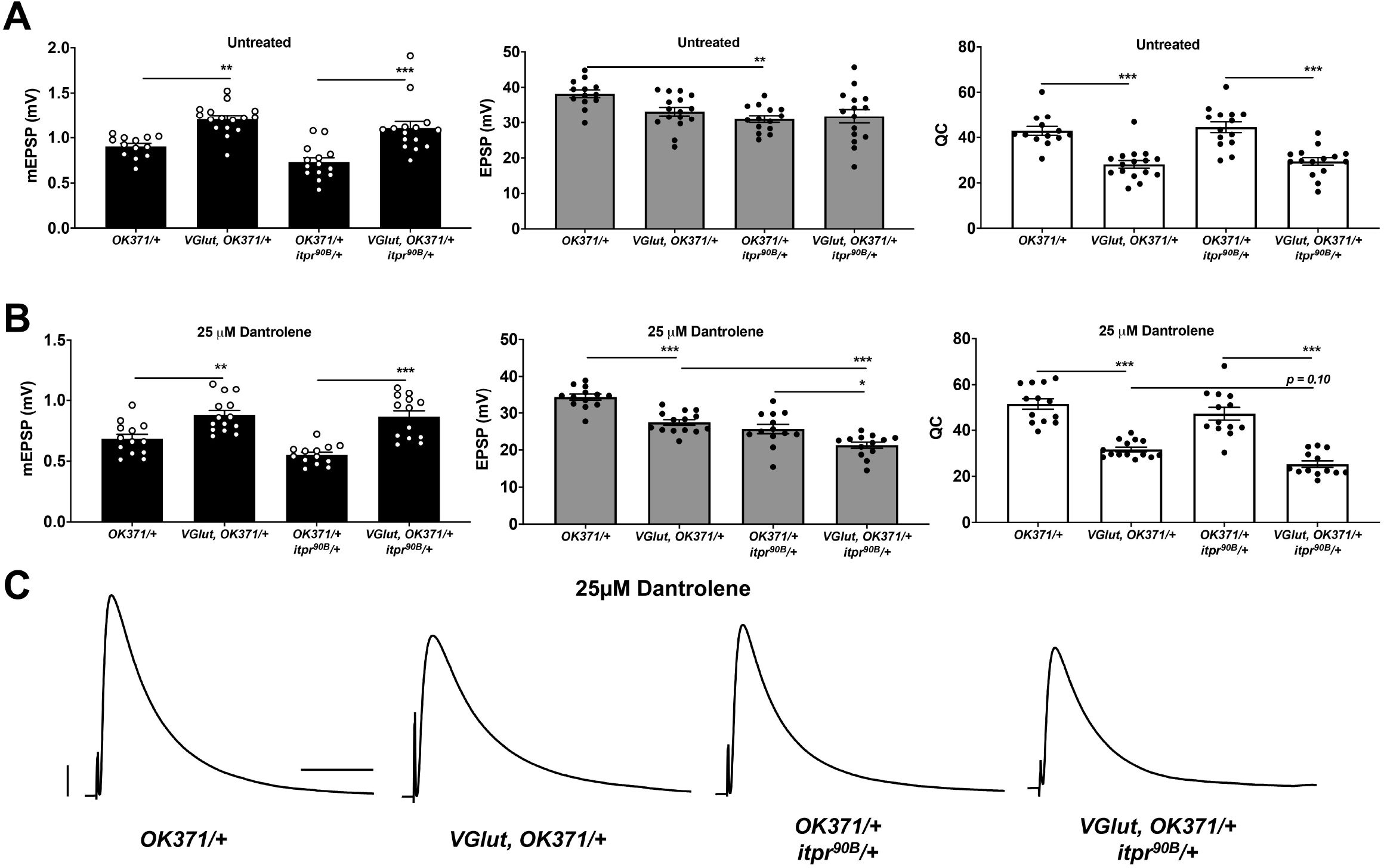
Genetic impairment of *itpr* combined with pharmacological impairment of RyR phenocopies prior genetic findings. Note: Untreated data for *OK371/+* and *VGlut, OK371/+* are repeated from Figure 1, for genetic background comparison. **(A)** Raw data for mEPSPs (left); raw data for EPSPs (middle); raw data for QC (right) for untreated genotypes as shown; bars are averages and error bars are ± SEM. **(B)** Data as in (A) but with 25 µM Dantrolene added to NMJ preps. * *p < 0*.*05, ** p < 0*.*01*, and *\*\*\* p < 0*.*001* by one-way ANOVA with Tukey’s post-hoc. **(C)** Representative EPSP traces. Scale bars are as in Figure 1.

In the absence of drug treatment, PHD proceeded normally (Fig. 5A). We noted that the untreated *OK371/+; itpr*^*90B*^*/+* heterozygous condition had a slightly diminished evoked amplitude compared to *OK371/+* (Fig. 5A, middle). Therefore, the *itpr*^*90B*^*/+* condition could be contributing some neurotransmission loss on its own. But in this case, the addition of *VGlut* transgenic expression to this heterozygous background did not further decrease evoked neurotransmission (Fig. 5A, middle), indicating normal PHP.

With 25 µM Dantrolene treatment, we noted that the mEPSP amplitudes were generally smaller than without treatment (compare Figs. 5A, B). Nevertheless, in the *VGlut-*overexpressing backgrounds, mEPSPs were still elevated compared to their respective controls (Fig. 5B). This indicated that in the presence of Dantrolene, there was still homeostatic pressure that could induce PHD. Additionally, with Dantrolene, EPSP amplitudes in *VGlut-*overexpressing lines were significantly decreased compared to their respective genetic controls (Figs. 5B, C). This was because of a marked decrease in QC (Fig. 5B). In particular, the *VGlut, OK371/+; itpr*^*90B*^*/+* condition (+ Dantrolene) had depressed evoked amplitudes compared either to the *VGlut, OK371/+* (+ Dantrolene) condition or to the *OK371/+; itpr*^*90B*^*/+* (+ Dantrolene) condition (Figs. 5B, C). Collectively, these data could indicate a cumulative neurotransmission defect when impairing both the IP_3_Rs and RyRs in a PHD-challenged background.

It is possible that strong impairment of RyRs could be sufficient to cause synthetic phenotypes in conjunction with the PHD regulation system. We ran additional pharmaco-genetic tests using a second sensitized genetic background, *OK371/RyR*^*E4340K*^ – both with and without drugs and with and without *UAS-VGlut* overexpression. Again, in the absence of pharmacological treatment, PHD proceeded normally in the heterozygous *OK371/RyR*^*E4340K*^ genetic background (Fig. 6A). With Dantrolene, mEPSPs became significantly larger when *VGlut* was expressed (Fig. 6B, left), but EPSPs were significantly reduced (Figs. 6B, middle, 6D) because of a decrease in quantal content (Fig. 6B, right).

**Figure 6.**
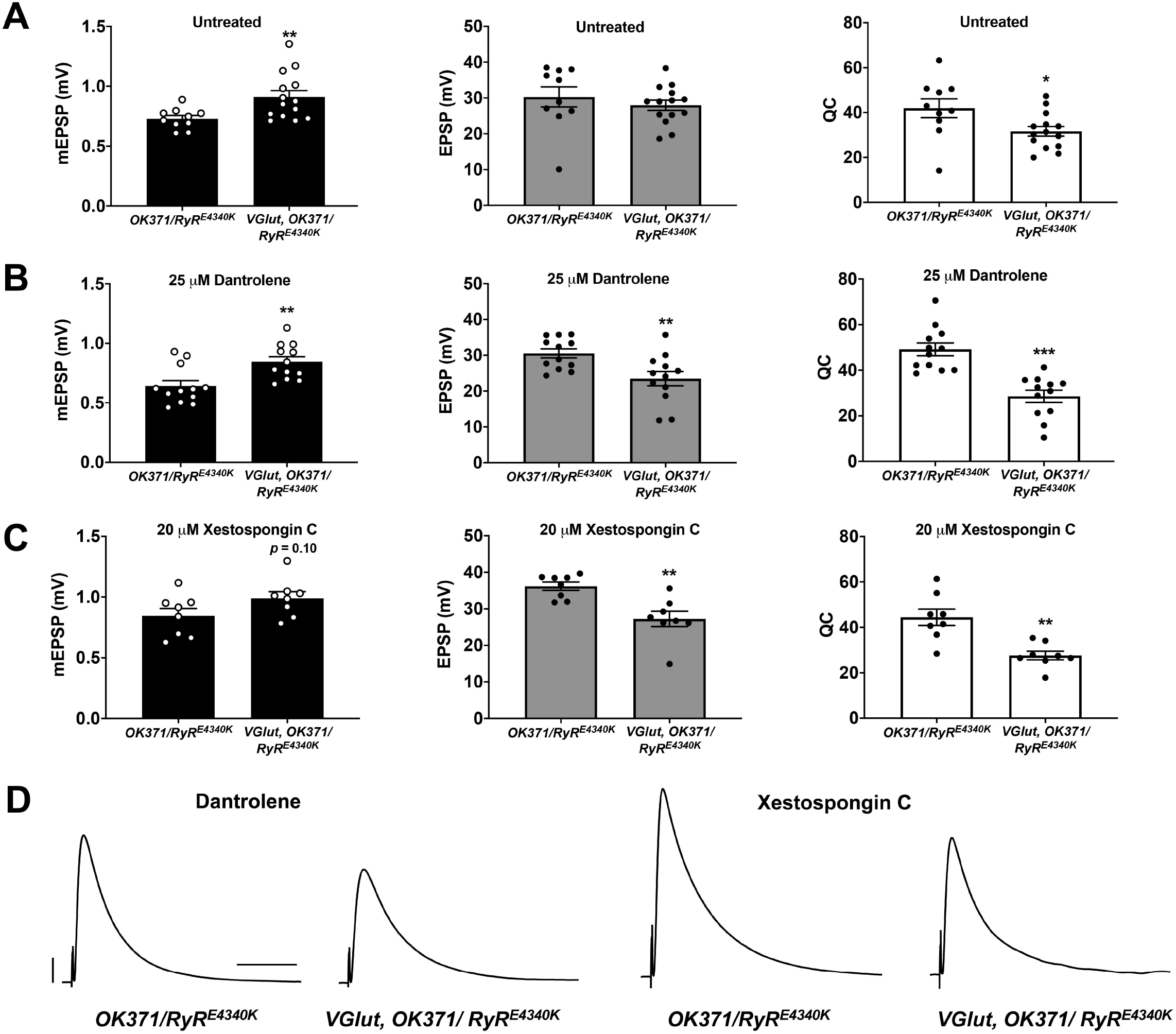
Additional pharmaco-genetic combinations phenocopy the genetic conditions. **(A)** Raw data for mEPSPs (left); raw data for EPSPs (middle); raw data for QC (right) for untreated genotypes as shown; bars are averages and error bars are ± SEM. **(B)** Data as in (A) but with 25 µM Dantrolene added to NMJ preps **(C)** Data as in (A) but with 20 µM Xestospongin C added to NMJ preps. **(D)** Representative EPSP traces. Scale bars are as in Figure 1. * *p < 0*.*05, ** p < 0*.*01*, and *\*\*\* p < 0*.*001* by Student’s T-Test comparing a control dataset (no VGlut overexpression) vs. an experimental dataset (VGlut overexpression).

Finally, we attempted the inverse pharmaco-genetic experiment from that in Figure 5. This time we used the IP_3_R inhibitor, Xestospongin C (Gafni et al., 1997; Wilcox et al., 1998) and the sensitized *OK371/RyR*^*E4340K*^ genetic background. We applied 20 μM Xestospongin C, both to *OK371/RyR*^*E4340K*^ NMJs and to *VGlut, OK371/RyR*^*E4340K*^ NMJs. mEPSPs were numerically larger when *VGlut* was overexpressed (Fig. 6C, left) – though interestingly, for the Xestospongin C dataset, the data did not achieve statistical significance for mEPSP size (*p* = 0.10, one-way ANOVA). This could indicate only weak to no homeostatic pressure in the presence of Xestospongin C. Nevertheless, EPSPs were significantly reduced (Figs. 6C, middle, 6D) because of a marked decrease in quantal content (Fig. 6C, right).

Taking all of these data together, for each case where we examined a dual impairment of RyR and IP_3_R the EPSP amplitudes were all quite low with concomitant VGlut overexpression (Figs. 3, 5, 6).

### PHD in very low extracellular calcium

We wondered how impairment of channels that mediate release of calcium from intracellular stores might cause the electrophysiological phenotypes that we observed. It could be the case that they are part of the PHD system. Or it could be the case that impairing these channels does not impinge upon PHD signaling itself – but their loss may sensitize the synapse to additional challenges, such as those brought on by PHD.

Our prior work suggested that these ER calcium store channels and the signaling systems that control them are required to maintain homeostatic potentiation throughout life (Brusich et al., 2015; James et al., 2019). We also found a related result: impairing Ca^2+^ store release mollified hyperexcitability phenotypes caused by gain-of-function Ca_V_2 amino-acid substitutions in the alpha1 subunit Cacophony. Ca_V_2 channels mediate synaptic calcium influx at the NMJ (Brusich et al., 2018). In light of these prior data, we considered two possibilities for PHD. One model is that the IP_3_R and RyR channels play a role in ensuring proper level of neurotransmission coincident with PHD. A different model is that calcium itself plays the important role. If this latter idea were true, it might be the case that lowering calcium influx into the presynaptic terminal would also be sufficient to interact with the PHD signaling process, ultimately lowering evoked transmission.

As a test, we measured release over a range of low extracellular calcium concentrations (0.2-0.5 mM). We examined six genotypes: 1) WT; 2) *w; OK371/+*; 3) *w; VGlut, OK371/+*; 4) *w; RyR*^*E4340K*^*/+; itpr*^*90B*^*/+*; 5) *w; OK371/RyR*^*E4340K*^; *itpr*^*90B*^*/+*; and 6) *w; VGlut, OK371/RyR*^*E4340K*^; *itpr*^*90B*^*/+*. To organize data and to calculate calcium cooperativity, we plotted quantal content as a function of calcium concentration, with the x-y axes on a log-log scale (Fig. 7A, B). To account for different Ca^2+^ driving forces in the different concentrations, we corrected QC for nonlinear summation in our plots and in our subsequent analyses (NLS Corrected QC) (Martin, 1955).

**Figure 7.**
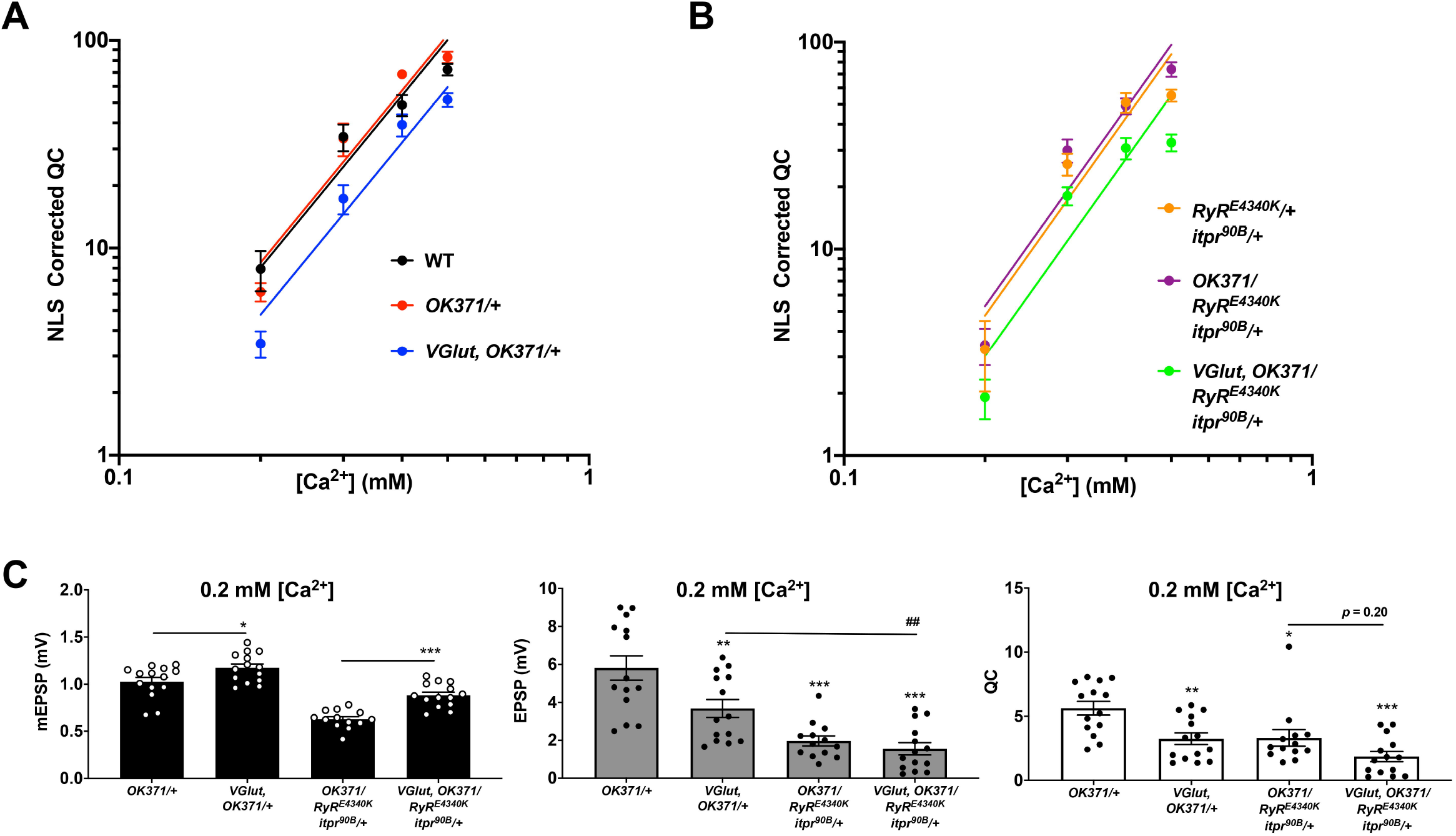
Ca^2+^ concentration-sensitivity of PHD execution. **(A)** Log-log plots of recording saline [Ca^2+^] vs. QC corrected for non-linear summation for WT, *OK371/+*, and *VGlut, OK371/+* conditions. Across the range of [Ca^2+^] examined, there is no significant difference in calcium cooperativity for these conditions (Nonlinear Regression, *p = 0*.*91*). **(B)** Data plotted as in (A) but this time with a double heterozygous *RyR*^*E4340K*^*/+; itpr*^*90B*^*/+* genetic background. Across the range of [Ca^2+^] examined, there is no significant difference in calcium cooperativity for these conditions (Nonlinear Regression, *p = 0*.*78*). **(C)** Raw data for mEPSPs (left); raw data for EPSPs (middle); raw data for QC (right). All data are for the indicated NMJ genotypes in 0.2 mM [Ca^2+^]; bars are averages and error bars are ± SEM. For mEPSPs, * *p < 0*.*05* and *\*\*\* p < 0*.*001* by Student’s T-Test, comparing PHD-challenged genotypes vs. unchallenged genetic controls. For EPSPs and QC, * *p < 0*.*05, ** p < 0*.*01*, and *\*\*\* p < 0*.*001* vs. *OK371/+*; *## p < 0*.*01*; EPSP and QC analyses done across multiple genotypes by one-way ANOVA with Tukey’s post-hoc.

Non-linear regression analyses revealed that there was no significant difference in calcium cooperativity between any of these genotypes over the range of extracellular [Ca^2+^] we tested (Fig. 7A, B). The calculated log-log slope values of the control PHD genotypes were: WT (log-log slope = 1.810), *w; OK371/+* (log-log slope = 1.884), and *w; VGlut, OK371/+* (log-log slope = 2.117). Comparing those three slopes with one another by nonlinear regression yielded no significant difference in slope (*p* = 0.91). The log-log slope values of the double heterozygous conditions were: *w; RyR*^*E4340K*^*/+*; *itpr*^*90B*^*/+* (log-log slope = 1.737), *w; OK371/RyR*^*E4340K*^; *itpr*^*90B*^*/+* (log-log slope = 2.102), and *w; VGlut, OK371/RyR*^*E4340K*^; *itpr*^*90B*^*/+* (log-log slope = 1.601). Comparing those slopes with one another also yielded no significant difference (*p* = 0.77).

Even though there was no significant difference in calcium cooperativity of release over the range of low [Ca^2+^] conditions examined, our data did show a very large drop in release between 0.3 and 0.2 mM [Ca^2+^] – specifically for the genotypes where PHD was induced by *UAS-VGlut* overexpression, or for the genotypes with a double heterozygous impairment of *RyR* and *itpr*. Examining the raw data at 0.2 mM [Ca^2+^], we observed that there was significant homeostatic pressure for PHD signified by mEPSP amplitude increases in the VGlut-overexpression background (Fig. 7C, left). Yet except for the control NMJs, EPSP amplitudes were very much diminished (Fig. 7C, middle) because of stark drops in QC (Fig. 7C, right).

Together, the data point to two conclusions. First, low extracellular calcium on its own appears to be a case where the synapse experiences a synergistic interaction with PHD challenge (Fig. 7C, *VGlut, OK371/+* data). Second, double heterozygous impairment of *RyR* and *itpr* appears to cause very low levels of baseline release in low calcium, irrespective of PHD challenge (Fig. 7C, middle; compare with Fig. 3C). Taken together, these data suggest that lowering presynaptic calcium by any means (impairing store release and/or impairing influx) is sufficient to impair evoked levels of excitation, in conjunction with a PHD challenge.

### PHD challenge interacts with impaired Ca_V_2 function

As a final test, we turned back to genetics. Drosophila Ca_V_2 channels mediate synaptic calcium influx at the NMJ. We used a hypomorphic mutant in the Ca_V_2 alpha1 subunit-encoding *cacophony* gene, *cac*^*S*^, to limit calcium influx. Ca_V_2 is essential for viability, but *cac*^*S*^ hypomorphs are viable and fertile (Kawasaki et al., 2000; Smith et al., 1998). Prior work showed that the *cac*^*S*^ homozygous condition dampens NMJ EPSP amplitude by about 70-80% (Frank et al., 2006); calcium imaging data suggest this is due to a ∼50% decrease in Ca^2+^ influx during evoked stimulation (Müller and Davis, 2012). Beyond this phenotype in baseline neurotransmission, *cac*^*S*^ hypomorphs also block PHP expression and PHP-associated increases in presynaptic calcium influx (Frank et al., 2006; Müller and Davis, 2012).

With a single cross, we generated hemizygous *cac*^*S*^*/Y; VGlut, OK371/+* male larvae (Fig. 8A). Compared to *cac*^*S*^*/Y* as a baseline mutant control, *cac*^*S*^*/Y; VGlut, OK371/+* NMJs have a marked increase in mEPSP size (Fig. 8B), indicating homeostatic pressure to induce PHD (Fig. 7B). However, comparing evoked potentials of those two conditions shows that *cac*^*S*^*/Y; VGlut, OK371/+* NMJs have much smaller EPSPs (Fig. 8C) and a very large decrease in QC (Fig. 8D).

**Figure 8.**
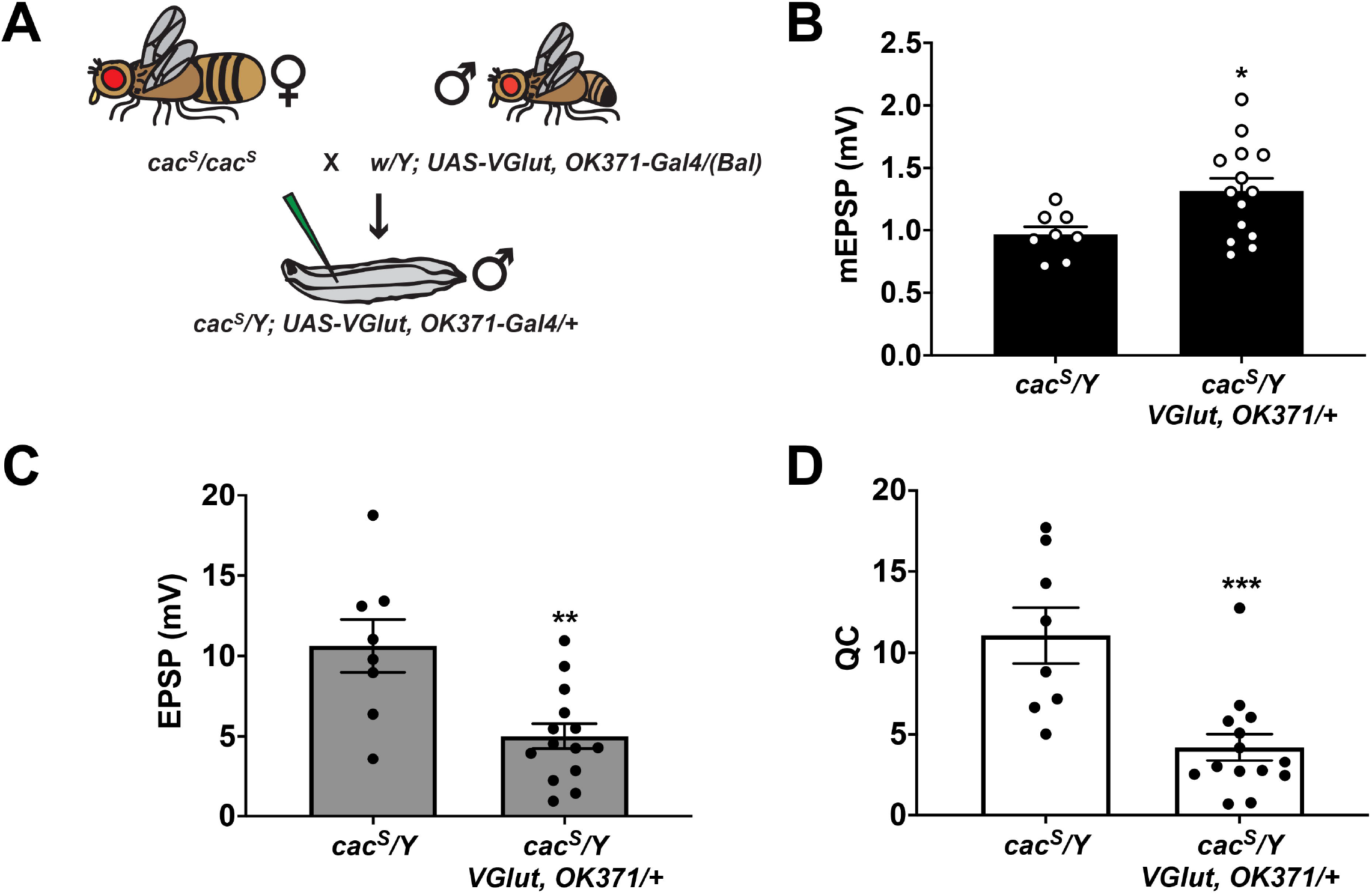
Partial impairment of Ca_V_2/Cacophony and PHD. **(A)** Crossing scheme for generating larvae for electrophysiological recording. Male larvae were hemizygous for the *cac*^*S*^ hypomorphic mutation. **(B)** Raw data for mEPSPs. **(C)** Raw data for EPSPs. **(D)** Raw data for QC. For (B)-(D), bars are averages and error bars are ± SEM. * *p < 0*.*05, ** p < 0*.*01*, and *\*\*\* p < 0*.*001* by Student’s T-Test comparing the control *cac*^*S*^ dataset (no *VGlut* overexpression) vs. the experimental *cac*^*S*^ dataset (*VGlut* overexpression).

## 4 Discussion

We began this study in search of genetic conditions that affect PHD (Fig. 2). While we did not find any conditions that result in a block of PHD, we did find conditions that provide insight into how calcium regulation may interact with this form of homeostatic plasticity to affect synapse function. When IP_3_R and RyR functions are partially impaired – either by genetics or by pharmacology – the NMJ still executes a PHD-like process. But that process goes beyond what is appropriate for the homeostatic pressure that is applied to the system. As a result, evoked potentials at the NMJ are much smaller than baseline (Figs. 3, 5, 6). A similar phenotype is observed when extracellular [Ca^2+^] is lowered to 0.2 mM (Fig. 7) and when the Ca_V_2 alpha1 subunit gene *cacophony* harbors a hypomorphic mutation, *cac*^*S*^ (Fig. 8).

This phenotype has important implications for proper control of synapse function. Taking our data together, we propose that perturbations that dampen calcium efflux from stores or perturbations that dampen calcium influx from the extracellular environment can both synergistically interact with a PHD challenge to control levels of evoked neurotransmission (Fig. 9).

**Figure 9.**
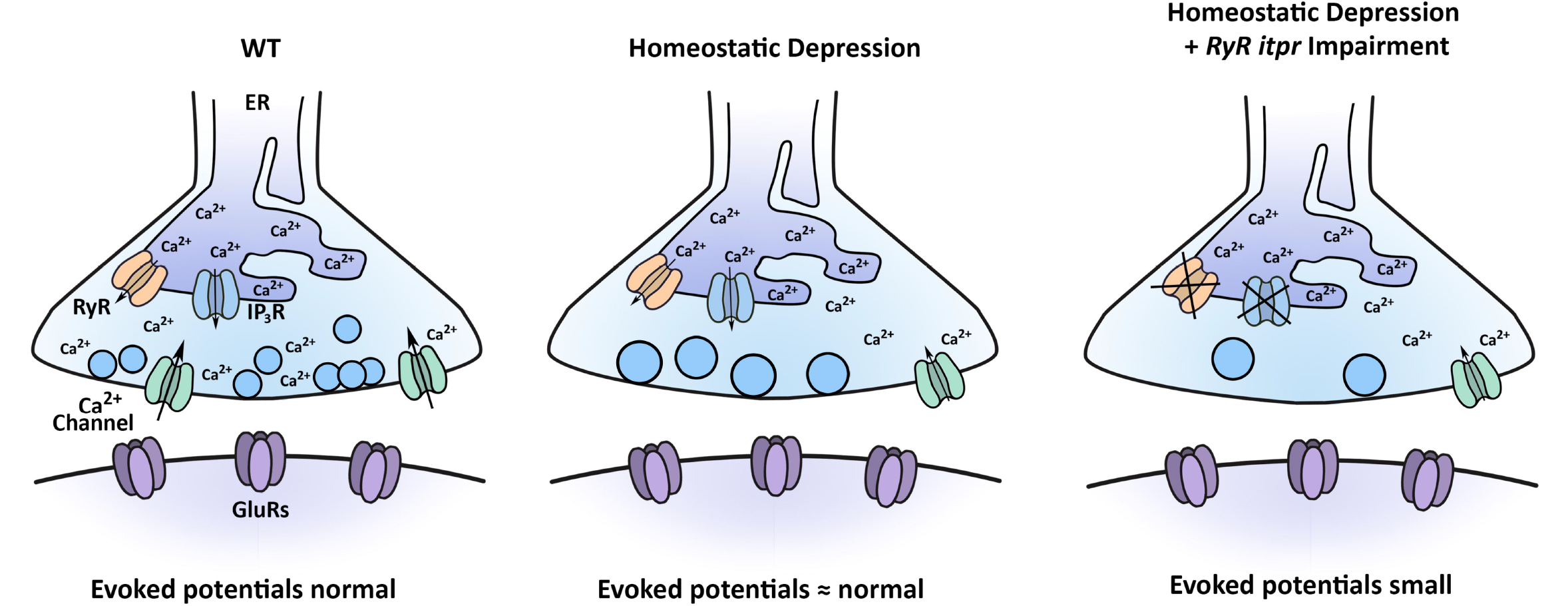
Model for how multiple calcium sources interact with the process of PHD. Under baseline conditions, Ca_V_2-type calcium channels contribute to synapse function, as may RyRs and IP_3_Rs. Under conditions inducing PHD, synaptic vesicles are enlarged, and QC is decreased, through regulation of sources of calcium. When PHD challenge is coupled with concomitant impairment of RyR and IP_3_R channels, evoked potentials are significantly diminished.

### Screen limitations

We did not identify conditions that blocked PHD, and here we discuss potential limitations of the screen. First, our primary assay was electrophysiology, and we employed a candidate-based method, similar to what has previously been documented in the field for PHP (Frank et al., 2020). By definition, candidate-based screens are limited in scope. Second, we focused on factors previously implicated in the maintenance of PHP function (or closely related signaling factors). The idea was that some factors needed to maintain synaptic homeostasis may be needed to orient the NMJ toward a proper, physiological level of function, regardless of the nature of the homeostatic challenge. This idea could have valence, but it was not guaranteed to produce mutant conditions with greater than normal evoked amplitudes in our screen.

Regarding the electrophysiological data, we did find instances in which the screened EPSP was numerically larger than the baseline for *VGlut, OK371/+*, but no instances identified as “PHD-blocking” (Fig. 2). The *VGlut, OK371/+* baseline evoked potential was high (∼35 mV), so it is possible that potential positives at a higher potential could be obscured by the limits of non-linear summation. There were also variations from line to line in resting membrane potential, input resistance, and the degree of mEPSP increase indicative of PHD challenge (Supplementary Table, S1). All of these parameters could contribute to false negatives for the screen. Unless a screen is done to saturation, there will be false negatives. It is important to interpret those parsimoniously. For our screen, we believe the way to interpret a negative is not to state that the screen definitively ruled out a factor – rather, the screen failed to rule in that factor for follow-up study.

### Similarities and differences with prior PHD studies at the NMJ

We were able to conduct a PHD screen using our recombinant stock with the *UAS-VGlut* and *OK371-Gal4* elements on the same chromosome. In principle, such a stock can pick up modifier mutations. The trade-off was a simplified, single-generation crossing scheme for genetic screens. Our recombinant stock with the driver and *UAS* elements in *cis* maintains consistent PHD challenge from generation to generation, and it behaves similarly electrophysiologically to *trans OK371/VGlut* combinations used in other studies (Daniels et al., 2004; Gaviño et al., 2015; Li et al., 2018).

There are differences between our study and the findings of other published work. Prior studies have used WT (or *w*^*1118*^) as a control background when compared to VGlut overexpression (Daniels et al., 2004; Gaviño et al., 2015; Li et al., 2018). This is a standard practice. Those studies reported precise PHD when comparing WT vs. *OK371/VGlut* third instar larvae – decreased QC at *OK371/VGlut* NMJs resulting in unchanged evoked transmission. We replicated this finding (Fig. 1). However, we also used our Gal4 driver stock background *OK371/+* as an additional control. For that comparison, we saw a slight depression in the evoked amplitude of *OK371, VGlut/+* NMJs (Fig. 1). One possibility is that our recombinant stock was acting as a sensitized background.

A second difference comes from the low extracellular calcium test. A low extracellular calcium experiment was previously done when VGlut overexpression was first characterized (Daniels et al., 2004). For that study, the authors showed that QC was significantly diminished compared to wild-type NMJs by the method of failure analysis. Taking the data of that study in aggregate, the authors concluded that PHD was intact in a variety of conditions, including saline with very low extracellular [Ca^2+^] (0.23 mM Ca^2+^, 20 mM Mg^2+^). Our study may appear to conflict with that study because we found that saline with very low [Ca^2+^] (0.2 mM Ca^2+^, 10mM Mg^2+^) is conducive an interaction with PHD, resulting in low evoked release. One possibility is that since the original study was examining failure percentage vs. WT – and not the absolute value of mEPSPs or EPSPs in low calcium, this might not be as easily observed. Other differences might be attributed genetic background or other differences in recording saline, like magnesium concentration.

Finally, one other study previously examined the effects of a *cac*^*S*^ mutation with concomitant VGlut overexpression (Gaviño et al., 2015). The authors did not find the low evoked potentials that we report. The major difference between that experiment and ours is that the prior work examined the *cac*^*S*^ mutation in an extracellular [Ca^2+^] (1.0 mM) that was double that of our study. The result was a Ca^2+^ driving force that yielded robust baseline EPSPs, even in the *cac*^*S*^ mutant background (Gaviño et al., 2015). Given our results with low calcium concentration (Fig. 7), a similar effect may be at work here.

### Known roles for calcium in controlling homeostatic plasticity

The notion that calcium contributes to successful homeostatic signaling is not new. Many roles for voltage-gated calcium channels in synaptic homeostasis are well-documented (Frank, 2014a, b). Prior to our study, there was evidence for voltage-gated calcium channel regulation for both NMJ PHP and PHD. For PHP, loss-of-function conditions in Ca_V_2/*cacophony* can impair or block this form of homeostatic regulation (Frank et al., 2006; Frank et al., 2009; Müller and Davis, 2012; Spring et al., 2016). Calcium imaging experiments suggest that the reason is because an increase in calcium influx through Ca_V_2 is required for the upregulation of quantal content during PHP, and mutant conditions like *cac*^*S*^ block this increase (Müller and Davis, 2012). Recent studies report that Cacophony and other active zone protein levels increase at the NMJ active zone in response to PHP homeostatic challenges (Böhme et al., 2019; Goel et al., 2019; Gratz et al., 2019). And work from mammalian systems mirrors these findings. For example, with mouse hippocampal cultures, TTX exposure induces a homeostatic decrease in presynaptic calcium influx (Zhao et al., 2011).

The converse appears true for PHD. Calcium imaging data from two different studies has shown a decrease in the size of calcium transients at the NMJ in response to presynaptic nerve firing in VGlut-overexpressing animals (Gaviño et al., 2015; Li et al., 2018). The data are mixed on how these decreased transients might come about during PHD. Using a tagged *UAS-cacophony* cDNA transgene, two studies verified that there was a reduction in the amount of GFP-tagged Cacophony alpha1 subunits in Ca_V_2 in a VGlut-overexpressing background (Gaviño et al., 2015; Gratz et al., 2019). However, one of these same studies demonstrated that if a tagged genomic construct is used instead, that same Ca_V_2 reduction is not observed (Gratz et al., 2019). Since the transgenic tagged Cacophony-GFP is the product of a single *cac* splice isoform (Kawasaki et al., 2002; Kawasaki et al., 2004), it could be the case that some isoforms are more dynamically trafficked at the synapse. Another possibility is that existing active zone components are somehow modulated during PHD. Regardless of the actual mechanism, the phenomenon appears conserved: again, with rodent hippocampal preparations, increased neuronal activity through gabazine exposure induces a PHD-like phenomenon ultimately resulting in decreases in calcium influx and release (Jeans et al., 2017; Zhao et al., 2011).

### How do calcium stores interact with PHD?

Calcium stores have been studied in the context of neurotransmission and plasticity. We know that endoplasmic reticulum (ER) can be visualized at Drosophila NMJ terminals (Summerville et al., 2016), and recently developed imaging tools employed in multiple systems (including at the Drosophila NMJ) show how nerve stimulation results in dynamic changes to ER lumenal calcium, (de Juan-Sanz et al., 2017; Handler et al., 2019; Oliva et al., 2020). In parallel, other groups working at the Drosophila NMJ have demonstrated important roles in baseline neurotransmission and in PHP for ER resident proteins (Genç et al., 2017; Kikuma et al., 2017). And from our prior work, we know that store calcium channels and upstream signaling components are important for maintaining the NMJ’s capacity for PHP throughout life (Brusich et al., 2015; James et al., 2019). We also know that disrupting these same factors can ameliorate hyperexcitability associated with gains of Ca_V_2 function (Brusich et al., 2018). Finally, from mammalian work it is clear that IP_3_Rs, RyRs, and intracellular calcium govern a variety of forms of neuroplasticity (Berridge, 2016), including paired pulse facilitation (Emptage et al., 2001), and modulation of voltage-gated calcium channel activity (Catterall, 2011; Lee et al., 2000).

If PHD were simply a matter of properly functioning neurotransmission machinery, then it is not entirely obvious why PHD would be so sensitive to the amount of calcium available such that evoked release would be impaired greatly either when store-operated channels were impaired or when the amount of influx was lowered. In our study, neurotransmission has not been lowered beyond a point of synapse failure. This means that there is still functional machinery. And PHD, per se, is not disrupted indeed, there is still depression.

With any type of homeostatic system, there not only needs to be error detection (large quantal size) and correction (decreased quantal content), but there also need to be brakes applied to the system to prevent some kind of overcorrection. At first glance, our data could suggest some manner of PHD “overcorrection.” In our view, this is an interesting and understudied type of phenomenon that could be examined in many homeostatic systems. But it is also true that the nature of the PHD challenge could simply represent a genetic background that renders the synapse sensitive to any additional insults.

So how exactly do levels of calcium (or the function of distinct types of calcium channels found at the synapse) ultimately affect excitation levels? This is a difficult problem. The first step might be to narrow the relevant tissue type(s) involved in PHD signaling. ER and store-operated channels are relevant to the functions of many tissues. In principle, our genetic loss-of-function manipulations to *itpr* and *RyR* could affect store-operated channels either in the neuron or in the muscle or in surrounding tissues like glia. Our pharmacological manipulations using Dantrolene and Xestospongin C could also affect multiple tissue types. Therefore, in principle, changing the levels of cytosolic calcium could either affect local signaling in the neuron, or it could result in aberrant signaling back to the presynaptic neuron, disorienting the homeostat.

We favor the idea that the relevant calcium signal is local in the motor neuron for two reasons. First, from our own data, we were able to observe the small evoked neurotransmission phenotype either with manipulations to store calcium or with manipulations that affect presynaptic calcium influx, including partial loss-of-function of neuronal *cacophony*. Second, a recent study puts forth data suggesting that when VGlut overexpression induces PHD, this happens exclusively because of excess presynaptic glutamate release, and presynaptic depression is initiated independent of any sort of postsynaptic response (Li et al., 2018). Such an autocrine signaling mechanism could very well reveal a role for intracellular calcium signaling in the presynapse.

## 8 Author Contributions

C.J.Y. and C.A.F. both did the following: designed research, performed research, analysed data, and wrote and edited the paper.

## 10 Acknowledgements

We thank members of the Frank lab for helpful comments. We thank the laboratories of Drs. Tina Tootle, Fang Lin, Toshihiro Kitamoto, Pamela Geyer, and Lori Wallrath for helpful discussions. We also thank Drs. Toshihiro Kitamoto, Joshua Weiner, Christopher Stipp, and Mark Stamnes for helpful feedback on an earlier written version of this study. An earlier version of this manuscript was released as a pre-print at *bioRxiv*, (Yeates and Frank, 2020).

## 11 Supplementary Material

Please see Supplementary Table S1 for raw data from the electrophysiology screen and a legend explaining the table.

## 12 Data Availability Statement

The raw data supporting the conclusions in this article will be made available by the authors, without undue reservation.

