## Supplemental Table S1 for "Homeostatic depression shows heightened sensitivity to synaptic calcium"

| Genotype | Condition | mEPSP (mV) | mEPSP Freq (Hz) | EPSP (mV) | QC | NLS QC | $I_R$ (MΩ) | RMP (mV) | n |
|---|---|---|---|---|---|---|---|---|---|
| *w*$^{1118}$ | 25°C | 0.9 ± 0.04 | 4.0 ± 0.5 | 34.1 ± 1.2 | 38.3 ± 2.0 | 72.4 ± 4.7 | 6.9 ± 0.5 | -64.7 ± 0.9 | 18 |
| *w; OK371/+* | 25°C | 0.9 ± 0.03 | 2.4 ± 0.2 | 38.2 ± 1.1 | 42.9 ± 2.0 | 83.0 ± 5.2 | 7.5 ± 0.5 | -70.7 ± 1.2 | 13 |
| *w; VGlut, OK371/+* | 25°C | 1.2 ± 0.04 | 4.8 ± 0.4 | 33.2 ± 1.2 | 28.1 ± 1.7 | 51.8 ± 4.0 | 5.8 ± 0.2 | -64.3 ± 1.0 | 16 |
| *> UAS-brp[RNA]*$^{JF01932}$*/+* | 25°C | 1.6 ± 0.14 | 4.4 ± 0.5 | 36.6 ± 1.7 | 24.5 ± 2.2 | 44.1 ± 4.5 | 10.4 ± 0.8 | -73.0 ± 1.8 | 7 |
| *> UAS-Calx[RNAi]*$^{JF02937}$*/+* | 25°C | 1.3 ± 0.12 | 5.1 ± 0.5 | 34.6 ± 2.9 | 28.2 ± 3.0 | 51.1 ± 7.1 | 6.9 ± 0.7 | -70.3 ± 1.6 | 9 |
| *> UAS-CaMKII*$^{Ala}$*/+* | 25°C | 1.1 ± 0.09 | 4.7 ± 0.2 | 31.8 ± 2.2 | 30.5 ± 2.6 | 51.1 ± 5.1 | 2.9 ± 0.3 | -69.9 ± 2.3 | 8 |
| *> UAS-CaMKII*$^{T287D}$*/+* | 25°C | 0.9 ± 0.05 | 6.3 ± 0.8 | 30.8 ± 3.9 | 32.4 ± 2.8 | 53.8 ± 7.7 | 1.8 ± 0.5 | -70.0 ± 3.5 | 4 |
| *> UAS-Cdc42*$^{N17}$*/+* | 25°C | 1.0 ± 0.09 | 4.8 ± 0.3 | 28.4 ± 2.2 | 29.0 ± 1.9 | 46.9 ± 4.3 | 5.6 ± 0.6 | -66.0 ± 1.0 | 5 |
| *w; VGlut, OK371/+; Csp*$^{EY22488}$*/+* | 25°C | 1.4 ± 0.11 | 5.4 ± 1.3 | 33.8 ± 2.2 | 25.2 ± 1.5 | 43.1 ± 3.5 | 5.9 ± 0.9 | -72.7 ± 3.9 | 6 |
| *> UAS-Csp[RNAi]*$^{GD10571}$*/+* | 25°C | 1.1 ± 0.05 | 4.5 ± 0.6 | 32.7 ± 1.5 | 30.5 ± 1.2 | 51.7 ± 3.0 | 4.4 ± 0.5 | -70.3 ± 1.0 | 7 |
| *> UAS-Csp[RNAi]*$^{KK109431}$*/+* | 25°C | 1.1 ± 0.05 | 4.8 ± 0.2 | 25.5 ± 2.4 | 23.0 ± 2.9 | 38.3 ± 7.5 | 5.5 ± 0.4 | -60.5 ± 0.3 | 7 |
| *> UAS-DnaJ-1[RNAi]*$^{HMS00688}$*/+* | 25°C | 1.3 ± 0.06 | 4.4 ± 0.3 | 33.6 ± 1.7 | 26.9 ± 1.3 | 47.3 ± 3.8 | 4.4 ± 0.7 | -71.1 ± 1.1 | 12 |
| *> UAS-DnaJ-1[RNAi]*$^{HMS00778}$*/+* | 25°C | 1.3 ± 0.19 | 6.2 ± 1.7 | 27.9 ± 3.2 | 23.2 ± 2.1 | 38.1 ± 5.0 | 7.9 ± 0.8 | -66.8 ± 1.5 | 10 |
| *> UAS-Eph[RNAi]*$^{GD2535}$*/+* | 25°C | 1.1 ± 0.09 | 4.3 ± 0.8 | 31.3 ± 3.0 | 30.0 ± 4.9 | 53.8 ± 11.0 | 5.4 ± 0.4 | -64.3 ± 2.9 | 6 |
| *> UAS-exn[RNAi]*$^{GD11079}$*/+* | 25°C | 1.2 ± 0.04 | 4.4 ± 0.5 | 32.9 ± 2.4 | 27.1 ± 1.3 | 47.7 ± 4.6 | 5.3 ± 0.5 | -68.2 ± 1.6 | 6 |
| *> UAS-Ephrin[RNAi]*$^{JF02365}$*/+* | 25°C | 1.3 ± 0.10 | 3.1 ± 0.3 | 29.1 ± 2.6 | 23.4 ± 2.2 | 38.8 ± 4.7 | 6.6 ± 0.5 | -65.1 ± 1.0 | 5 |
| *> UAS-Gαq[RNAi]*$^{JF02464}$*/+* | 25°C | 1.0 ± 0.05 | 3.1 ± 0.4 | 32.5 ± 1.0 | 32.1 ± 1.7 | 58.6 ± 3.8 | 5.6 ± 0.4 | -62.2 ± 0.7 | 7 |
| *w; VGlut, OK371/Gαq*$^{28}$ | 25°C | 1.0 ± 0.04 | 6.4 ± 0.2 | 31.5 ± 1.1 | 31.9 ± 2.1 | 57.3 ± 4.8 | 4.6 ± 0.3 | -62.3 ± 0.5 | 10 |
| *> UAS-htl[RNAi]*$^{GD14457}$*/+* | 25°C | 1.0 ± 0.06 | 4.6 ± 0.6 | 28.8 ± 1.2 | 28.9 ± 1.9 | 48.9 ± 3.8 | 4.7 ± 0.3 | -61.4 ± 0.6 | 8 |
| *> UAS-IP*$_3$*-sponge*$^{m30}$*/+* | 25°C | 1.2 ± 0.10 | 4.3 ± 0.4 | 28.3 ± 1.9 | 25.1 ± 2.1 | 39.8 ± 4.3 | 6.6 ± 0.3 | -70.3 ± 1.4 | 8 |
| *> UAS-IP*$_3$*-sponge*$^{m49}$*/+* | 25°C | 1.1 ± 0.07 | 6.0 ± 0.7 | 33.9 ± 1.5 | 30.9 ± 2.9 | 55.4 ± 6.6 | 3.7 ± 0.5 | -68.3 ± 1.7 | 6 |
| *w; VGlut, OK371/+; itpr*$^{90B}$*/+* | 25°C | 1.1 ± 0.07 | 6.5 ± 0.4 | 31.8 ± 1.9 | 29.4 ± 1.7 | 50.1 ± 3.8 | 3.4 ± 0.5 | -69.7 ± 1.5 | 15 |
| *w; VGlut, OK371/+; itpr*$^{ka1091}$*/+* | 25°C | 1.1 ± 0.04 | 5.5 ± 0.6 | 28.4 ± 1.3 | 26.6 ± 1.1 | 42.3 ± 2.7 | 5.5 ± 0.3 | -68.5 ± 1.6 | 12 |
| *w; VGlut, OK371/+; itpr*$^{ug3}$*/+* | 25°C | 1.1 ± 0.08 | 5.8 ± 0.4 | 29.5 ± 1.3 | 27.3 ± 1.4 | 45.2 ± 2.6 | 5.1 ± 0.4 | -65.7 ± 1.1 | 19 |
| *> UAS-itpr[RNAi]*$^{GLC01786}$*/+* | 25°C | 1.0 ± 0.05 | 4.2 ± 0.3 | 25.6 ± 2.3 | 26.7 ± 2.9 | 43.1 ± 6.2 | 5.7 ± 0.3 | -65.1 ± 1.0 | 13 |
| *> UAS-itpr[RNAi]*$^{HMC03351}$*/+* | 25°C | 1.0 ± 0.05 | 4.1 ± 0.3 | 29.2 ± 1.4 | 31.3 ± 2.1 | 53.3 ± 4.5 | 6.9 ± 0.5 | -62.5 ± 0.7 | 10 |
| *> UAS-Mad[RNAi]*$^{JF01958}$*/+* | 25°C | 1.0 ± 0.04 | 4.0 ± 0.4 | 28.3 ± 2.7 | 28.1 ± 2.8 | 45.3 ± 6.2 | 5.3 ± 0.5 | -68.4 ± 1.3 | 7 |
| *> UAS-mGluR[RNAi]*$^{HMS00191}$*/+* | 25°C | 1.1 ± 0.09 | 3.8 ± 0.4 | 32.1 ± 0.6 | 29.5 ± 2.1 | 51.3 ± 3.7 | 6.0 ± 0.5 | -65.6 ± 0.6 | 10 |
| *> UAS-mGluR[RNAi]*$^{JF01958}$*/+* | 25°C | 1.2 ± 0.07 | 3.3 ± 0.2 | 30.1 ± 2.3 | 25.6 ± 2.5 | 43.8 ± 5.3 | 7.5 ± 0.4 | -68.0 ± 1.2 | 11 |
| *> UAS-mGluR[RNAi]*$^{KK102270}$*/+* | 25°C | 1.0 ± 0.06 | 4.3 ± 0.5 | 36.0 ± 1.9 | 38.1 ± 2.6 | 73.2 ± 6.4 | 6.8 ± 0.4 | -66.2 ± 1.6 | 9 |
| *> UAS-Plc21C[RNAi]*$^{HMS00600}$*/+* | 25°C | 1.0 ± 0.06 | 2.9 ± 0.2 | 30.2 ± 1.7 | 31.1 ± 1.0 | 52.8 ± 2.7 | 5.1 ± 0.3 | -64.7 ± 0.9 | 11 |
| *> UAS-Rbp[RNAi]*$^{JF02471}$*/+* | 25°C | 1.0 ± 0.04 | 3.4 ± 0.3 | 28.1 ± 1.9 | 27.6 ± 1.5 | 45.3 ± 3.7 | 5.3 ± 0.5 | -64.0 ± 0.8 | 8 |
| *w; VGlut, OK371/RyR*$^{16}$ | 25°C | 1.0 ± 0.05 | 5.4 ± 0.4 | 34.5 ± 1.2 | 34.7 ± 1.9 | 64.5 ± 4.4 | 5.5 ± 0.5 | -66.6 ± 1.0 | 18 |
| *w; VGlut, OK371/RyR*$^{E4340K}$ | 25°C | 0.9 ± 0.05 | 3.8 ± 0.3 | 28.0 ± 1.4 | 31.7 ± 2.2 | 51.2 ± 4.3 | 6.3 ± 0.6 | -65.3 ± 1.2 | 14 |
| *> UAS-RyR[RNAi]*$^{HM05130}$*/+* | 25°C | 1.2 ± 0.1 | 3.7 ± 0.6 | 34.2 ± 2.5 | 29.2 ± 2.8 | 53.9 ± 6.8 | 6.0 ± 0.5 | -66.3 ± 1.6 | 6 |
| *w; VGlut, OK371/RyR*$^{E4340K}$*; itpr*$^{90B}$*/+* | 25°C | 1.1 ± 0.06 | 5.0 ± 0.8 | 23.3 ± 1.2 | 22.1 ± 1.6 | 32.7 ± 3.0 | 5.4 ± 0.2 | -67.0 ± 0.9 | 19 |
| *w; VGlut, OK371/+ ; Smn*$^{73Ao}$*/+* | 25°C | 1.2 ± 0.18 | 6.1 ± 0.7 | 31.6 ± 2.1 | 29.4 ± 3.3 | 51.0 ± 7.2 | 1.6 ± 0.2 | -69.2 ± 1.0 | 10 |
| *w; VGlut, OK371/+ ; Smn*$^{f05960}$*/+* | 25°C | 1.2 ± 0.06 | 4.8 ± 0.7 | 36.8 ± 1.1 | 31.4 ± 1.6 | 58.9 ± 3.2 | 5.7 ± 0.5 | -69.3 ± 0.8 | 9 |
| *> UAS-Smn[RNAi]*$^{GL00581}$*/+* | 25°C | 1.2 ± 0.06 | 5.1 ± 0.3 | 28.3 ± 2.1 | 23.5 ± 1.9 | 38.1 ± 4.2 | 6.1 ± 0.5 | -69.0 ± 1.3 | 12 |
| *> UAS-Smn[RNAi]*$^{HMC03832}$*/+* | 25°C | 1.3 ± 0.1 | 5.1 ± 0.5 | 31.1 ± 2.2 | 23.7 ± 1.3 | 39.7 ± 3.3 | 7.3 ± 0.5 | -68.9 ± 1.7 | 8 |
| *> UAS-Smn[RNAi]*$^{JF02057}$*/+* | 25°C | 1.0 ± 0.07 | 3.6 ± 0.2 | 20.7 ± 1.7 | 21.5 ± 1.3 | 29.6 ± 2.3 | 5.7 ± 0.3 | -68.3 ± 1.9 | 8 |
| *> UAS-Snap25[RNAi]*$^{HMS01367}$*/+* | 25°C | 1.2 ± 0.06 | 2.5 ± 0.2 | 37.4 ± 3.0 | 31.9 ± 2.9 | 60.9 ± 7.7 | 3.1 ± 0.7 | -71.7 ± 1.3 | 7 |
| *> UAS-Snap25[RNAi]*$^{JF02615}$*/+* | 25°C | 1.1 ± 0.14 | 2.9 ± 0.6 | 26.1 ± 2.7 | 24.0 ± 2.9 | 37.8 ± 5.7 | 5.5 ± 0.7 | -65.5 ± 1.2 | 7 |
| *> UAS-Snapin[RNAi]*$^{JF02692}$*/+* | 25°C | 1.4 ± 0.13 | 5.2 ± 0.6 | 35.3 ± 2.2 | 26.0 ± 2.1 | 46.0 ± 3.8 | 3.3 ± 0.5 | -71.6 ± 1.8 | 6 |
| *> UAS-wit[RNAi]*$^{10776R-1}$*/+* | 25°C | 1.2 ± 0.06 | 3.8 ± 0.7 | 37.0 ± 1.1 | 31.1 ± 1.3 | 60.7 ± 2.7 | 2.8 ± 0.4 | -66.4 ± 1.2 | 8 |

**Supplemental Table S1.** Raw electrophysiology data for the genetic screen depicted in Fig. 2. Due to limited space, the genotypes are depicted by shorthand explained here. ">" denotes the data for a cross in which virgin females of the genotype *w; CyO-GFP/UAS-VGlut, OK371-Gal4* are crossed to males harboring a *UAS*-driven transgene or RNAi line. ">" implies a genetic background of *w/+; UAS-VGlut, OK371-Gal4/+* inherited from the mother in order to drive (>) the heterozygous UAS condition inherited from the father. For the cases where mutations to an endogenous locus were screened, the entire genotype is written out. Average values ± SEM are given for each parameter. n = number of NMJs for the genotype. Parameters include miniature excitatory postsynaptic potential (mEPSP) amplitude, miniature frequency (mEPSP Frq), excitatory postsynaptic potential (EPSP) amplitude, quantal content (QC), QC corrected for non-linear summation (NLS QC), input resistance ($I_R$), and resting membrane potential (RMP). Gray shading divides the data by gene targeted.
